# Shared genetics and couple-associated environment are major contributors to the risk of both clinical and self-declared depression

**DOI:** 10.1101/076398

**Authors:** Yanni Zeng, Pau Navarro, Charley Xia, Carmen Amador, Ana M. Fernandez-Pujals, Pippa A. Thomson, Archie Campbell, Reka Nagy, Toni-Kim Clarke, Jonathan D. Hafferty, Blair H. Smith, Lynne J. Hocking, Sandosh Padmanabhan, Caroline Hayward, Donald J. MacIntyre, David J Porteous, Chris S. Haley, Andrew M. McIntosh

## Abstract

**Background:** Both genetic and environmental contributions to risk of depression have been identified, but estimates of their effects are limited. Commonalities between major depressive disorder (MDD) and self-declared depression (SDD) are also unclear. Dissecting the genetic and environmental contributions to these traits and their correlation would inform the design and interpretation of genetic studies.

**Methods:** Using data from a large Scottish family-based cohort (GS:SFHS, N=21,387), we estimated the genetic and environmental contributions to MDD and SDD. Genetic effects associated with common genome-wide genetic variants (SNP heritability) and additional pedigree-associated genetic variation and Non-genetic effects associated with common environments were estimated using linear mixed modeling (LMM).

**Findings:** Both MDD and SDD had significant contributions from effects of common genetic variants, the additional genetic effect of the pedigree and the common environmental effect shared by couples. The correlation between SDD and MDD was high (r=1⋅00, se=0⋅21) for common-variant-associated genetic effects and moderate for both the additional genetic effect of the pedigree (r=0⋅58, se=0⋅08) and the couple-shared environmental effect (r=0⋅53, se=0⋅22).

**Interpretation:** Both genetics and couple-shared environmental effects were the major factors influencing liability to depression. SDD may provide a scalable alternative to MDD in studies seeking to identify common risk variants. Rarer variants and environmental effects may however differ substantially according to different definitions of depression.

**Funding:** Study supported by Wellcome Trust Strategic Award 104036/Z/14/Z. GS:SFHS funded by the Scottish Government Health Department, Chief Scientist Office, number CZD/16/6.

## Introduction

Depression has a pattern of familial aggregation, which implies the influence of genetic effects, common environmental effects shared by relatives, or both. The genetic component (heritability) has been estimated by a twin study of major depressive disorder (MDD) to be 37%^1^. The SNP heritability (heritability attributed to common genetic variants) of MDD varies across populations and samples (0⋅21~0⋅32)^2,3^. Subsequently, a ‘children of twins’ study found a significantly greater risk of depression in children of depressed monozygotic (MZ) twins than in the offspring of their non-depressed twin^4^. This implies a potential environmental effect of parental depression on offspring^4^. Studies have also shown that having a partner with psychiatric disorder may increase an individual's risk of MDD^5,6^, but meta-analytic studies suggest no effect of the shared sibling environment and other studies have postulated more complex relationships^7^. Whilst each of these studies separately provided evidence for the genetic and familial environmental components in depression, a precise separation of these potential effects should involve estimating them simultaneously in the same model and has yet to be achieved.

The accurate separation and estimation of the genetic and environment components on liability to depression provides crucial information, as it reveals the upper limit of the genetic effects, the likely returns of genetic studies and the potential for accurate risk predictions^8,9^. Genetic studies attempting to map causal variants have been performed for various definitions of depression. These include clinically assessed depression, self-report of clinical diagnosis of depression and self-reported depressive symptoms^10–13^, but the findings are generally inconsistent, which suggests intrinsic heterogeneity across depression definitions. This is further supported by the fact that studies show very different heritability estimates of depression phenotypes in addition to MDD 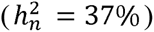^1^: perceived stress: 44% ^14^; nine depression definitions in women: 21%-45%^15^; depressive symptom scores in childhood: 79%^16^ and depressive symptoms in an elderly population: 69% in women and 64% in men^17^.

Because of the heterogeneity across depression definitions, there has been a long debate about the correct phenotype for depression genetic studies. Studies using clinically-assessed depression definitions could provide findings that are directly informative for clinical application. However, the resources required for data collection are generally very high^10^. As an alternative, measuring self-reported depression requires fewer resources and is rapidly becoming available for many population-based datasets^12,13^. To date, the largest published GWAS of major depression has yielded 15 significant loci for a self-reported clinical diagnosis^12^.

Given that important progress is being made from GWASs on different depression definitions, it becomes increasingly important to understand the similarities and dissimilarities across different definitions. Particularly, the difference in genetic and environmental loadings might underpin the inconsistent results from genetic studies across different depression definitions. Therefore, dissecting the phenotypic variance of each depression definition and understanding the similarities and dissimilarities between those phenotypes in the context of both genetic and common environmental components is particularly important for interpreting the results from published depression studies and for informing genetically relevant depression phenotypes for future studies.

In this study we sought to partition the phenotypic variation of the diagnosed depression (MDD) and the self-declared depression (SDD) into its genetic and familial environment components using Linear Mixed Modeling (LMM). Specifically, we utilized data from Generation Scotland: Scottish Family Health Study (GS:SFHS), a large Scottish cohort with extensive family relationship information and genome-wide genotype data to answer two questions. First, when simultaneously considering multiple genetic effects and familial environmental effects in the model, what are the major contributions to variation in MDD and SDD, respectively? Second, what is the contribution of each of the identified major contributing components to the overall correlation between MDD and SDD?

## Methods and Materials

The Tayside Research Ethics Committee (reference 05/S1401/89) provided ethical approval for the study.

## Datasets

### Generation Scotland: Scottish Family Health Study (GS:SFHS)

contains 21,387 subjects (N_male_=8,772, N_female_=12,615; Age_mean_=47⋅2), who were recruited from the registers of collaborating general practices. At least one first-degree relative aged 18 or over was required to be identified for each participant^18^. Genotyping data were generated using the Illumina Human OmniExpressExome -8-v1.0 array^19^. Details of genotyping are described in detail elsewhere^18^. Outlier individuals were removed from the sample^20^. Quality control (QC) of genotyped SNPs used inclusion thresholds: missing SNPs per individual ≤2%, SNP genotype call rate ≥98%,minor allele frequency (MAF) >1% and Hardy-Weinberg equilibrium P value > 1x10^−6^. In total, 561,125 genotyped autosomal SNPs passed QC criteria and were available for 19,994 participants.

#### Phenotypes

Lifetime Diagnosis of MDD: The Structured Clinical Interview for DSM-IV(SCID) was used^21^: participants who screened positive (21⋅7%) were invited to continue to an interview using the SCID modules for mood disorders^22^. Participants who screened positive but refused to undergo the structured clinical interview (N=507, 2⋅4%) and those with a diagnosis of bipolar disorder (N=76) were excluded from the study.

Self-declared depression (SDD): the participants were invited to answer the following question “please mark an X in the box if you have been affected by depression”.

### Partitioning the phenotypic variation

Based on the framework of the Genomic-relationship-matrix restricted maximum likelihood (GREML) method, Xia *et al*. (2016) developed a method to estimate 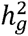 (additive genetic effect contributed by common variants, namely SNP heritability), 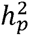 (additional additive genetic effect contributed by pedigree associated variation), 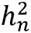 (total additive genetic effect, namely narrow-sense heritability) and a number of familial environmental components simultaneously^23^. This is performed by fitting variance-covariance matrices representing common genetic effects, pedigree-related-genetic effects, and recent and early family environmental effects simultaneously in the mixed linear model^23^, building on previous work by Zaitlen *et al*^24^. This approach enables estimation of the contribution of each genetic and family environmental component and here we applied it to MDD and SDD.

In detail, for each trait, two genomic relationship matrices, ***G*** (genomic relationship matrix) and ***K*** (kinship matrix created by modifying ***G***)^23,24^ and three environment relationship matrices, ***F*** (environmental matrix representing nuclear-family-member relationships), ***S*** (environmental matrix representing full-siblings relationships) and ***C*** (environmental matrix representing couple relationships) (Figure 1, Text s1)^23^, were fitted separately or simultaneously in a LMM (Text s2). The correspondent variance component 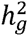 (common variants-associated genetic effect, represented in ***G***), 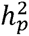(additional genetic effect of pedigree, represented in ***K***), 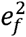 (environmental effect from nuclear family, represented in ***F***), 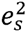(environmental effect from full sibling relationship, represented in ***S***) and 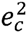 (environmental effect from couple relationship, represented in ***C***) were estimated and tested using LRT and Wald tests (Text s2). To simplify the model description, the following codes were used to represent the matrices fitted in the models:-e.g. ‘***GKFSC***’ was the full model which fits all five matrices as random effects simultaneously, and ‘***GKC***’ represents the model where the genomic relationship matrix, the kinship matrix and the environmental matrix representing couple relationships were simultaneously fitted.

**Figure 1.**
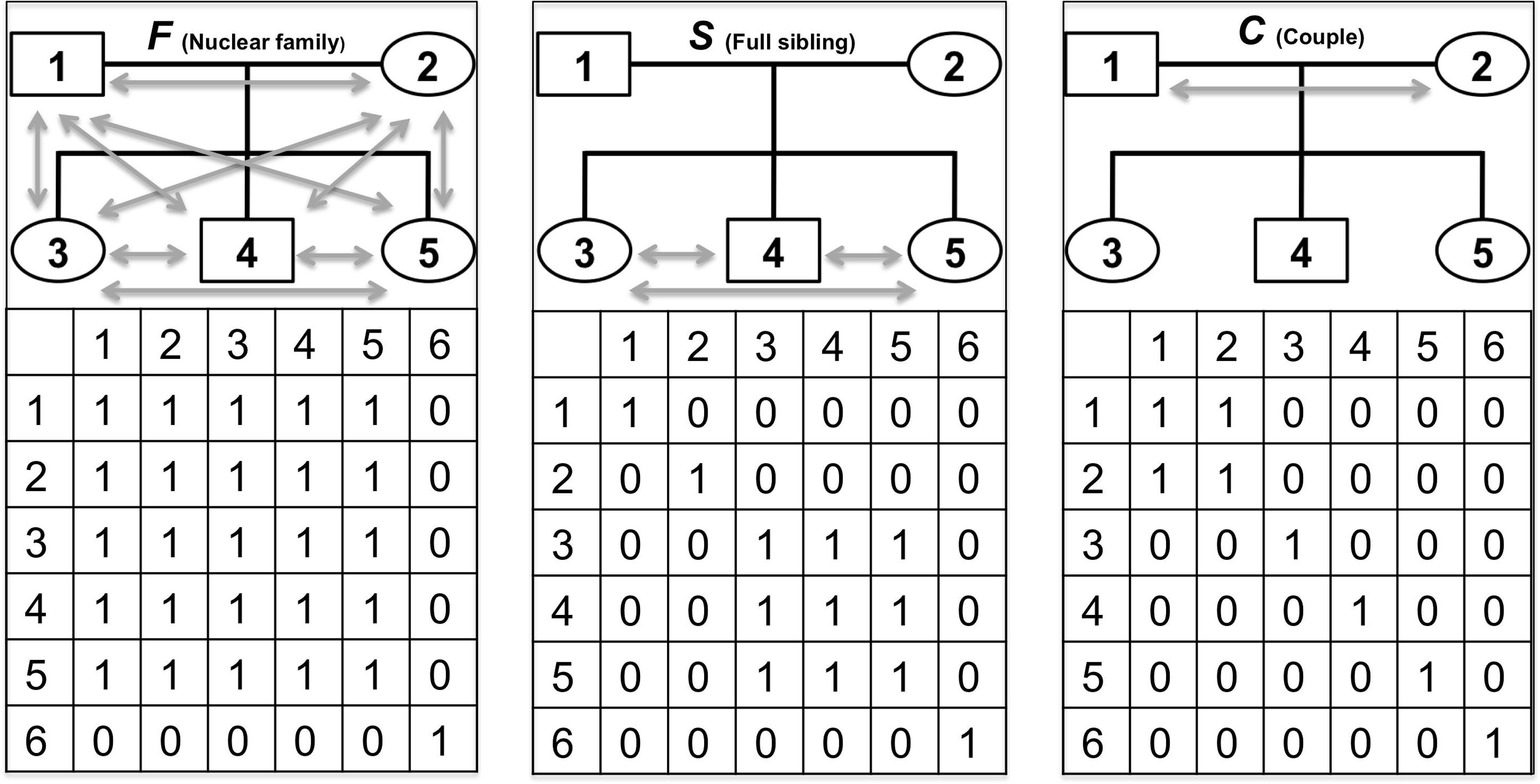
The design of the environment relationship matrices.

#### Stepwise model selection for identifying major contributing components

##### Backward stepwise selection^23^

The selection started with the full model ‘***GKFSC***’. LRT and Wald tests were conducted for each variance component. Variance component was removed from the model if (1) it failed to obtain significance (5%) in both tests and (2) among the variance components satisfying (1), it had the highest P value in the Wald test. This process was performed repeatedly until all the remaining components obtained significance in at least one test.

##### Forward stepwise selection

apart from backward selection, we additionally implemented Forward stepwise approach for best model selection. In this case the selection started with a null model (without fitting any matrices). Matrices were then added into the model one at a time. LRT and Wald tests were conducted for each variance component. A variance component was added if (1) it obtained significance (5%) in both tests, (2) the adding of this component did not lead to the variance components already in the model becoming non-significant in both tests and (3) among the variance components satisfying both (1) and (2), it had the lowest P value in the Wald test. This process was repeated until no more variance components satisfied criteria (1), (2) and (3).

#### Bivariate GREML analysis for MDD and SDD

Using the results obtained from the above analyses, we then estimated the relative contributions of each major contributing genetic and environmental components to the correlation between MDD and SDD. We used the GCTA-bivariate GREML analysis^25,26^ to estimate the correlation between the two traits in the SNP-associated genetic component, the pedigree-associated genetic component and the shared couple environment component simultaneously. Each estimated correlation was tested to determine significance of its difference from both 0 and 1.

## Results

In GS:SFHS, among the 19,994 participants with genome-wide genotyping data, we recognized 1,742 pairs of couples, 8,458 pairs of full siblings and 20,019 pairs of members living in the same nuclear family. In this dataset, 99⋅5% (19,896/19,994) participants have MDD diagnosis information (2,659 MDD cases and 17,237 controls) and 98% (19,603/19,994) participants have answered the question for self-reported depression (1,940 SDD cases and 17,633 controls).

A full model was first utilized to partition the phenotypic variation of each trait into five potentially influential sources: the additive genetic effect contributed by common variants 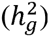, the additional additive genetic effect associated with pedigree 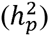, the environmental effect shared between nuclear family members 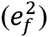, the environmental effect shared between couples 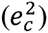 and the environmental effect shared between full siblings 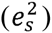 (Figure 1). The results were presented in Table 1. For MDD, 10% (se=5%) of the phenotypic variance is attributable to the common genetic variants 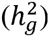 and 20% (se=12%) is to the additional genetic variation associated with pedigree 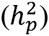. For SDD, 22% (se=7%) of the phenotypic variance is attributable to the common genetic variants 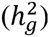 and 50% (se=15%) is to the additional genetic variation associated with pedigree 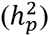. The proportion of total additive genetic determinant (narrow-sense heritability: 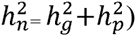) was 30% for MDD and 72% for SDD. For the three familial environmental components, LRT for the couple-shared environmental effect was significant for SDD (0⋅17 (se=0⋅10)), but was not significant for MDD (although it obtained a non-zero point estimate: 0.03 (se=0.09)). The environmental effects shared between nuclear family members and full-siblings were not significant for either trait. Compared with the a reduced model that does not account for environmental effects (the ***GK*** model), the full model obtains lower estimates of the genetic components for both traits, suggesting that the full model effectively reduced the confounding environmental effects when estimating the heritability (Figure 2,Table 1).

**Figure 2.**
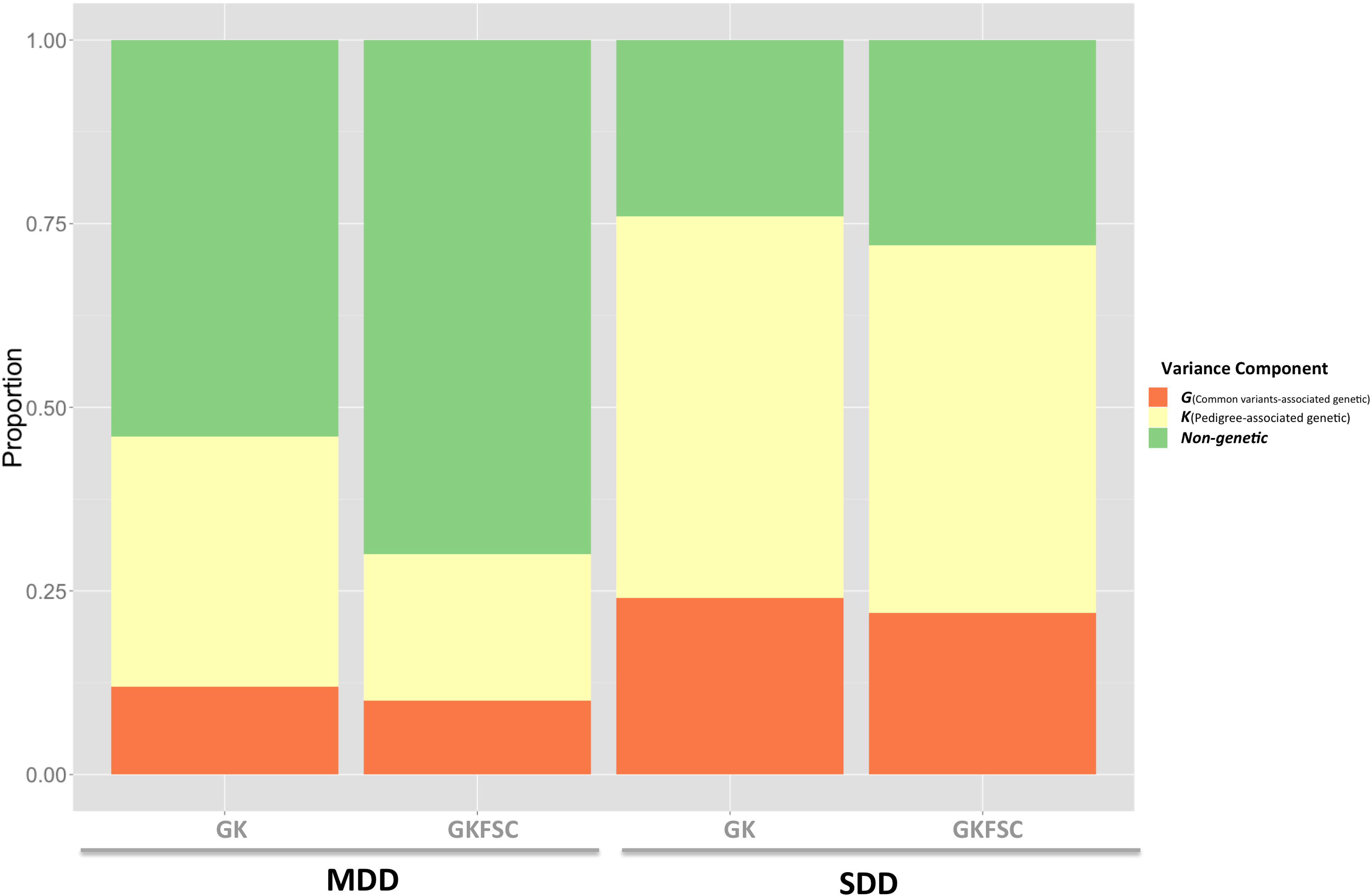
Comparison of the proportion of variance explained by the common-variants- and pedigree-associated genetic components estimated in the model that only account for the two genetic components (the ***GK*** model) and in the full model that accounts for two genetics and three shared-environmental effects (the ***GKFSC*** model). SDD: self declared depression, MDD: major depressive disorder.

**Table 1.** Variance component analyses results for MDD and SDD. Variance component analyses were performed on MDD and SDD using the genetic model (***GK***), the model accounting for all of the two genetic and three familial environmental effects (the full model ***GKFSC***), and the models selected by forward or backward selection. NS: the variance component was not significant in LRT test.

|  |  |  | **G**<br>(Common variants<br>-associated genetic) | **K**<br>(Pedigree-associated<br>genetic) | **F**<br>(Nuclear family) | **S**<br>(Full sibling) | **C**<br>(Couple) |
| --- | --- | --- | --- | --- | --- | --- | --- |
| **Trait** | **Description** | **Model** | $h_g^2(\text{se})$ | $h_{kin}^2(\text{se})$ | $e_f^2(\text{se})$ | $e_s^2(\text{se})$ | $e_c^2(\text{se})$ |
| **MDD** | Genetics only | **GK** | 0.12(0.05) | 0.34(0.06) |  |  |  |
|  | Full | **GKFSC** | 0.10(0.05) | 0.20(0.12) | 0.09(0.06)<sup>ns</sup> | 0.00(0.04)<sup>ns</sup> | 0.03(0.09)<sup>ns</sup> |
|  | Forward selection | **GKC** | 0.12(0.05) | 0.35(0.06) |  |  | 0.14(0.07) |
|  | Backward selection | **GF** | 0.15(0.04) |  | 0.16(0.03) |  |  |
| **SDD** | Genetics only | **GK** | 0.24(0.06) | 0.52(0.07) |  |  |  |
|  | Full | **GKFSC** | 0.22(0.06) | 0.50(0.15) | 0.04(0.07)<sup>ns</sup> | 0.00(0.04)<sup>ns</sup> | 0.17(0.10) |
|  | Forward selection | **GKC** | 0.24(0.06) | 0.53(0.07) |  |  | 0.22(0.07) |
|  | Backward selection | **GKC** | 0.24(0.06) | 0.53(0.07) |  |  | 0.22(0.07) |

As shown in previous work, the full model, although accounts for all of the five potentially influential effects, may have difficulty in separating major contributors to depression from minor contributors because of correlated effects^23^. In order to address this problem^23^, we applied stepwise model selection^23^ to identify the major contributors to variation in the two depression traits.

Using forward stepwise selection, the common variant-associated-genetic effect, pedigree-associated-genetic effect and couple-shared environmental effects were retained in the final model for both MDD and SDD (the ***GKC*** model as shown in Table s1 and Table 1). Using backward stepwise selection, for MDD, only the common variant-associated-genetic component and shared-nuclear-family component were retained in the final model (the ***GF*** model as shown in Table s2A). For SDD, common variant-associated-genetic, pedigree-associated-genetic effect and couple-shared environmental effects was selected (the ***GKC*** model as shown in Table s2B). The relative contribution of each variance component to SDD and MDD in the ***GKC*** model is shown in Figure 3.

**Figure 3.**
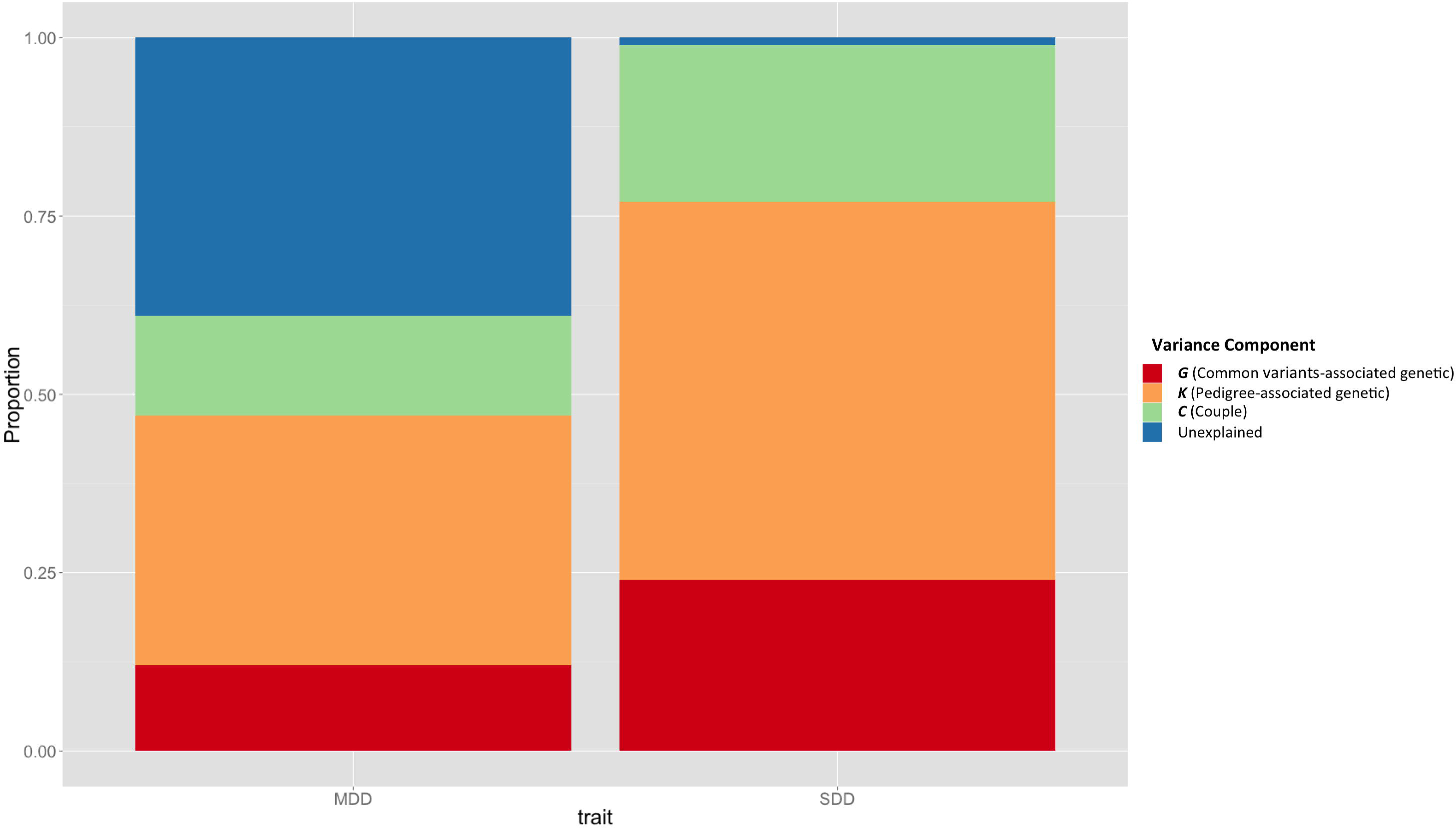
Sources of phenotypic variance and the proportion of variance they explained in the most parsimonious model (***GKC***) for both SDD and MDD. SDD: self declared depression, MDD: major depressive disorder.

The model that included common variants- and pedigree-associated-genetic and couple-shared environmental effects (the ***GKC*** model) was selected as the most parsimonious model in three out of the four model selections for the two traits (for the interpretation of ***GF*** model for MDD see Text s3). The three effects were therefore recognized as the major variance contributors for depression. We further explored the contribution of each of these contributors to the correlation between MDD and SDD. The phenotypic correlation between MDD and SDD is 0⋅45 (Phi coefficient, *P* < 2⋅2x10^−16^). The estimate of the common variants-associated genetic correlation was 1⋅00 (se=0⋅21), with this estimate being significantly different from 0 (*P*_*lrt_H0:r=0*_= 2⋅8x10^−5^) and not significantly different from 1(*P*_*lrt_H0:r=1*_=0⋅5). The estimate of the pedigree-associated genetic correlation was 0⋅58 (se=0⋅08, *P*_*lrt_H0:r=0*_=2⋅1x10^−10^, *P*_*lrt_H0:r=1*_=5⋅6x10^−7^), and the estimate of the couple-shared environmental correlation was 0⋅53 (se=0⋅22, *P*_*lrt_H0:r=0*_= 0⋅06, *P*_*lrt_H0:r=1*_ = 0⋅05)(Table 2).

**Table 2.** Bivariate GCTA-GREML estimates of the correlation of each variance component between SDD and MDD using ***GKC*** model.

|  | Test | MDD | SDD |
| --- | --- | --- | --- |
| ***G***<br>(Common variants-associated genetic) | $h_g^2$ | 0.12(0.05) | 0.24(0.06) |
| | $r_g$ | 1.00(0.21) | |
| ***K***<br>(Pedigree-associated genetic) | $h_{kin}^2$ | 0.35(0.06) | 0.53(0.07) |
| | $r_{kin}$ | 0.58(0.08) | |
| ***C***<br>(Couple) | $e_c^2$ | 0.14(0.07) | 0.22(0.07) |
| | $r_c$ | 0.53(0.22) | |

## Discussion

This study presented the first application of variance component models, so far as any of the authors are aware, simultaneously accounting for five genetic and familial environment factors to MDD and SDD. We made use of a large Scottish sample composed of both close and distant relatives with genome-wide genotyping data in a series of variance components analyses. We showed that the common variants- and pedigree-associated genetics and the couple-shared environmental effects are the major factors influencing liability to both clinically assessed and self-reported depression. After breaking down the correlation between the two traits into each contributing effect, the correlation between SDD and MDD was very high (r=1⋅00, se=0⋅21) for common variant-associated genetic effects and moderate for both the pedigree-associated genetic effect(r=0⋅58, se=0⋅08) and the couple-shared environmental effect (r=0⋅53, se=0⋅22).

The estimates of both the common variant-associated and the pedigree associated genetic effects are much lower in MDD 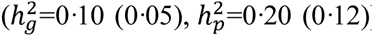 compared with those in SDD 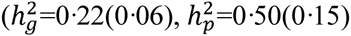, suggesting the difference in terms of the loading of the genetic components in the clinical and self-reported depression definitions. The point estimate of SNP heritability 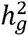 for MDD 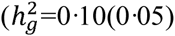 is lower than that from a mega-analysis for 9 cohorts of European ancestry (0⋅21 (0⋅02)^3^ and this may be due to the intrinsic heterogeneity of MDD^11^. The pedigree-associated genetic component 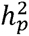 measures the additional genetic effects co-segregating in the pedigree, such as the effect from rare and structural variants. Notably, using the full model which partitions the narrow sense heritability into the common-variant-associated component 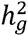 and the pedigree-associated component 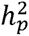, and accounts for multiple familial environmental effects in the meantime, the estimated 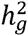 accounted for 31% of 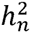 in both MDD and SDD, which is lower than previous estimates for other complex traits suggesting more than 50% of the 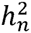 is accounted by 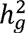 ^24^. On the contrary, the 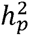 accounted for more than two third of the 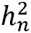 suggesting an important role for rare and structural variants in both clinical- and self-declared depression.

We failed to detect a full-sibling environmental effect in either trait, which was consistent with a previous study that suggested minimum contribution of the sibling environmental effect in MDD ^1^. Our findings suggested that the two genetic effects and the couple-shared environmental effects were the most significant contributors of depression. In total, the selected ***GKC*** model explained 60% (se=8%) and 98% (se=9%) of the phenotypic variation in MDD and SDD, respectively (Figure 3). Strikingly, the effect of the shared couple environment contributed as much as 14% (se=7%) to 22% (se=7%) of the phenotypic variance in MDD and SDD respectively. The role of the couple effect has been suggested in a Finnish study which showed that the partner’s MDD status associates with the MDD risk in non-psychiatric subjects^5^. Our results provided additional evidence for the couple-shared environmental effect and indicated that the magnitude of this effect is high. These results suggest that the inclusion of partners, whilst logistically attractive for researchers of depression, might introduce confounding if additional adjustment is not made. It may be helpful for future genetic studies to be aware of these potential biases and to either avoid the recruitment of couples, or model their effects appropriately. This finding also suggests that in clinical practices, apart from the traditional attention to the family history of patients, the mental health status of the spouse should also been treated as an important indication of risk.

The couple-shared environmental effect detected in depression traits could be confounded by the non-random mating in those phenotypes. For example, assortative mating has been observed in multiple psychiatric disorders including MDD^27^. When assortative mating exists in the trait of interest, the 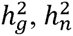 and 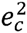 may be estimated with bias if this effect is not accounted for appropriately^23,28^. On the other hand, ignoring the couple-shared environment effect in the study of assortative mating may also lead to inaccurate measurement of the degree of assortation. In GS:SFHS, the mean onset age of MDD was 31⋅7 years^29^, suggesting a possibility of the participants developed depression before marrying, and therefore a possibility of assortative mating. Whilst the average age of couples in GS:SFHS is 59, suggesting that most couples have been living in the same household for a long time, therefore providing an opportunity for development of a couple-shared environment effect. Studies providing the age of marriage with large enough sample sizes to separate young and old couples would benefit the discrimination of the assortative mating effect and the couple-shared environment effect.

Finally, we estimated the genetic and environment correlation between MDD and SDD. The results revealed a very high correlation in the common variants-associated genetic component, a moderate correlation in the pedigree-associated genetic component and in the couple-shared environment component. This suggests that there are strong genetic similarities between the depression phenotypes amongst common variants. In contrast, pedigree-associated genetic variation and shared environment may underpin important differences between these traits. This has important implications for the design of future molecular studies of depression. According to our results, SDD could be a good candidate depression phenotype for MDD in population-based studies which target the genetic effects attributable to common variants. However, for family-based studies where the targeted genetic effect is from rare variants, the lower genetic correlation may impede the replication of findings from MDD in SDD.

## Acknowledgements

We are grateful to the families who took part in GS:SFHS, the GPs and Scottish School of Primary Care for their help in recruiting them, and the whole GS team, which includes academic researchers, clinic staff, laboratory technicians, clerical workers, IT staff, statisticians and research managers.

## Authors’ contributions

Conceived and designed the experiments: YNZ AMM CSH PN DJP SPH BHS LJH AC. Performed the experiments: CH AC RN DJM AMFP. Analyzed the data: YNZ. Contributed reagents/materials/analysis tools: CX CSH PN YNZ AMM CA AMFP TKC. Collecting samples: BHS LJH SP DJM DJP AMM CSH. Wrote the paper: YNZ CSH AMM PN CA TKC JDH PAT and all the other authors. Raised funding: AMM CSH DJP.

## References

1. Sullivan PF, Neale MC, Kendler KS. Genetic epidemiology of major depression: review and meta-analysis. Am J Psychiatry 2000; 157(10): 1552–62.

2. Lubke GH, Hottenga JJ, Walters R, et al. Estimating the genetic variance of major depressive disorder due to all single nucleotide polymorphisms. Biol Psychiatry 2012; 72(8): 707–9.

3. Lee SH, Ripke S, Neale BM, et al. Genetic relationship between five psychiatric disorders estimated from genome-wide SNPs. Nature genetics 2013; 45(9): 984-+.

4. Singh AL, D’Onofrio BM, Slutske WS, et al. Parental depression and offspring psychopathology: a children of twins study. Psychological medicine 2011; 41(7): 1385–95.

5. Joutsenniemi K, Moustgaard H, Koskinen S, Ripatti S, Martikainen P. Psychiatric comorbidity in couples: a longitudinal study of 202,959 married and cohabiting individuals. Soc Psych Psych Epid 2011; 46(7): 623–33.

6. Desai S, Schimmack U, Jidkova S, Bracke P. Spousal similarity in depression: a dyadic latent panel analysis of the panel study of Belgian households. Journal of abnormal psychology 2012; 121(2): 309–14.

7. Olino TM, Lewinsohn PM, Klein DN. Sibling similarity for MDD: evidence for shared familial factors. Journal of affective disorders 2006; 94(1-3): 211–8.

8. Makowsky R, Pajewski NM, Klimentidis YC, et al. Beyond missing heritability: prediction of complex traits. PLoS genetics 2011; 7(4): e1002051.

9. Tenesa A, Haley CS. The heritability of human disease: estimation, uses and abuses. Nature Reviews Genetics 2013; 14(2): 139–49.

10. consortium C. Sparse whole-genome sequencing identifies two loci for major depressive disorder. Nature 2015; 523(7562): 588–91.

11. Major Depressive Disorder Working Group of the Psychiatric GC, Ripke S, Wray NR, et al. A mega-analysis of genome-wide association studies for major depressive disorder. Molecular psychiatry 2013; 18(4): 497–511.

12. Hyde CL, Nagle MW, Tian C, et al. Identification of 15 genetic loci associated with risk of major depression in individuals of European descent. Nature genetics 2016; advance online publication.

13. Okbay A, Baselmans BML, De Neve J-E, et al. Genetic variants associated with subjective well-being, depressive symptoms, and neuroticism identified through genome-wide analyses. Nature genetics 2016; 48(6): 624–33.

14. Bogdan R, Pizzagalli DA. The heritability of hedonic capacity and perceived stress: a twin study evaluation of candidate depressive phenotypes. Psychological medicine 2009; 39(2): 211–8.

15. Kendler KS, Neale MC, Kessler RC, Heath AC, Eaves LJ. A population-based twin study of major depression in women. The impac t of varying definitions of illness. Archives of general psychiatry 1992; 49(4): 257–66.

16. Thapar A, Mcguffin P. A Twin Study of Depressive Symptoms in Childhood. Brit J Psychiat 1994; 165: 259–65.

17. McGue M, Christensen K. The heritability of depression symptoms in elderly Danish twins: Occasion-specific versus general effects. Behav Genet 2003; 33(2): 83–93.

18. Smith BH, Campbell H, Blackwood D, et al. Generation Scotland: the Scottish Family Health Study; a new resource for researching genes and heritability. BMC medical genetics 2006; 7: 74.

19. Gunderson KL. Whole-genome genotyping on bead arrays. Methods in molecular biology 2009; 529: 197–213.

20. Amador C, Huffman J, Trochet H, et al. Recent genomic heritage in Scotland. Bmc Genomics 2015; 16.

21. First MB, Spitzer RL, Gibbon M, Williams JB. Structured Clinical Interview for DSM-IV^®^ Axis I Disorders (SCID-I), Clinician Version, Administration Booklet: American Psychiatric Pub; 2012.

22. First MB, Spitzer RL, Gibbon M, Williams JB. Structured clinical interview for DSM-IV-TR axis I disorders, research version, patient edition: SCID-I/P, 2002.

23. Xia C, Amador C, Huffman J, et al. Pedigree- and SNP-Associated Genetics and Recent Environment are the Major Contributors to Anthropometric and Cardiometabolic Trait Variation. PLoS genetics 2016; 12(2): e1005804.

24. Zaitlen N, Kraft P, Patterson N, et al. Using Extended Genealogy to Estimate Components of Heritability for 23 Quantitative and Dichotomous Traits. PLoS genetics 2013; 9(5).

25. Yang J, Lee SH, Goddard ME, Visscher PM. GCTA: a tool for genome-wide complex trait analysis. American journal of human genetics 2011; 88(1): 76–82.

26. Lee S H, Yang J, Goddard ME, Visscher PM, Wray NR. Estimation of pleiotropy between complex diseases using single-nucleotide polymorphism-derived genomic relationships and restricted maximum likelihood. Bioinformatics 2012; 28(19): 2540–2.

27. Nordsletten AE, Larsson H, Crowley JJ, Almqvist C, Lichtenstein P, Mataix-Cols D. Patterns of Nonrandom Mating Within and Across 11 Major Psychiatric Disorders. Jama Psychiat 2016.

28. Keller MC, Garver-Apgar CE, Wright MJ, et al. The genetic correlation between height and IQ: shared genes or assortative mating? PLoS genetics 2013; 9(4): e1003451.

29. Fernandez-Pujals AM, Adams MJ, Thomson P, et al. Epidemiology and Heritability of Major Depressive Disorder, Stratified by Age of Onset, Sex, and Illness Course in Generation Scotland: Scottish Family Health Study (GS:SFHS). PloS one 2015; 10(11): e0142197.

